# The cardiac STAT3 intercalated disc specific expression in tail suspension rat

**DOI:** 10.1101/2021.06.20.449202

**Authors:** He Dongyu, Hu Aihua, Tong Jun, Zheng Chang, Liu Yiming, Chen Zhibin, Gao Lingfeng, Wang Yang

**Affiliations:** Bachelor of medicine and surgery curriculum, Hainan Medical University; Morphology laboratory, Hainan Medical University; Hainan province key laboratory of brain science research and transformation Extreme environment sports medicine laboratory

**Keywords:** Microgravity simulation, Right ventricle remodeling, Intercalated disc

## Abstract

**Background:** The cardiovascular system is significantly agitated by loss of gravity. In microgravity, the body fluids shift toward the thoracic cavity, induced the heart becomes more spherical. This further increased the cardiac preload with an increasing of transmural central venous pressure, affects the right heart ventricles to tolerating the enhanced preload on the right ventricular wall.

**Method:** In this study we investigated the rat right ventricle remodeling in simulating persistent microgravity by using tail-suspension model, examined the remolding of the heart and the specific STAT3 expression in right heart myocardium.

**Result:** The results indicated that microgravity induced heart remodeling included a significant increasing of the ventricular weight in the left. However, the right ventricle was not increased significantly in the microgravity simulation rats. The histological study demonstrated that the outstanding development on right ventricular wall which included the gap junction remodeling and STAT3 signaling protein specific accumulation in the right ventricles.

**Conclusion:** The results existed that the right cardiac ventricle has a distinctive remodeling process during microgravity simulation which was not the muscular hypertrophy and relative weight increasing, but manifested the STAT3 accumulation and the electrical gap junction remodeling. The effect of microgravity induced right ventricle remodeling and the STAT3 specific accumulation can be used for multi-purpose research.

## Introduction

The cardiovascular system is significantly agitated by loss of gravity. When the physiological hydrostatic gradients are vanished in exposure to microgravity, blood translocate from the lower part of the body towards the chest virtually doubling the amount of blood within the heart. The heart shape responds very quickly.to the bulk redistribution of fluid shift. In parabolic flight, blood and fluids shift along the body axis toward the cranium during the simulation of zero-G (1), echocardiography showed that the heart becomes more spherical. Body liquids redistribution excites a number of consistent physiological reactions as well as a cascade of secondary adaptation processes. The reactions occur in central venous pressure and size of the heart cavities. Microgravity is the condition in which objects appear to be weightless. The effects of microgravity can be seen when astronauts and objects float in space. Microgravity induced the heart changes its shape from an oval (like a water-filled balloon) to a round ball (an air filled balloon) (2). The short-term spaceflights cause a shift of ∼2 liters of blood and fluids from the lower body segments to the upper part of the body, that goes along with an increase of cardiac output of ∼20% (3). The observed increase of cardiac output is thought to be due to increased venous return. This further increased the cardiac preload with an increasing of transmural central venous pressure. Such hemodynamic feature affects the right heart ventricles to tolerating the enhanced preload on the right ventricular wall.

In this study we investigated the rat right ventricle remodeling in simulating persistent microgravity by using tail-suspension model, examined the specific expression of STAT3 in intercalated disc in enhanced preload cardiomyopathy caused body fluid redistribution.

## Methods

### Animal simulation model

A total of 10 SPF WiStar male rat, 4 weeks old, were purchased from the Experimental Animal Center of Hainan medical university. The rats were assigned to two groups: a control group (n=5), and microgravity simulation group (n=5). The study protocol was approved by the Ethics Committee of Hainan medical university, and all animal experiments were performed in accordance with the guidelines of the animal care and use committee of Hainan medical university.

Rats were microgravity simulation at a two-week tail suspension. During the simulation, rats were caged in an air-conditioned space at 23°C with 12-h light/12-h dark cycle. The tail was suspended and adjusted to prevent the hindlimbs from touching the cage surface. The body was tilted about 30°, allowing the forelimbs to maintain contact with the cage floor. The rat moves in a circular area to gain access to food and water. The control rats were housed in the cages with a tail-lift harness without suspension. Rats were sacrificed after 2 weeks of simulation.

### Cardiac right ventricle myocardium preparation

Rat hearts were harvested under the binocular dissecting microscope. The cardiac surface epicardium layer, and the intracardiac layers that included trabeculae, endocardium, were removed. The mid-wall of the ventricular myocardium was prepared in *Ringer*’s solution. The ventricles were fixed in 4% paraformaldehyde overnight at room temperature, then dehydrated in ethanol series, embedded in paraffin, and sectioned into 7-μm slices. The non-specific staining induced by endogenous peroxidase activity, endogenous alkaline phosphatase (AP) activity, and non-immunological binding of the specific immune sera were blocked by 3% bovine serum albumin (Thermo Fisher Scientific Inc.). The indirect method was introduced to immunohistological staining. The unlabeled anti-STAT3 primary antibody (unconjugated rabbit polyclonal to STAT3, ab226942, Abcam), 1:1000 diluted with PBS buffer solution, incubated in humidity chamber overnight at 4°C. The HRP labeled goat anti rabbit IgG secondary antibody (AB_228332, Invitrogen), 1:3000 diluted with PBS buffer solution, incubated 1h at room temperature. The slides finally colored with diaminobenzidine (DAB). The cellular nucleus was stained with Hematoxylin (Mayer’s Hematoxylin Solution, Sigma-Aldrich).

### The quantities analysis

Before paraformaldehyde fixation, the wet ventricular weight was measure compared between the control and simulation group.

The ventricle morphology was studied by using immunohistochemical staining slide. The ventricular myocardium images were capture by upright microscope (Nikon eclipse 80i, Nikon instruments Inc., Tokyo, Japan) with NIS elements imaging software (Nikon Instruments Inc. NY, U.S.A.). The images were computer-assistant imaging analyzed by NIHimage (ImageJ 1.53a, National institutes of health, U.S.A.). The intercalated disc and the STAT3 positive area pixels were extracted by using the adjust mode in ImageJ 1.53a (Wayne Rasband, NIH, U.S.A.).

### Statistics

The mean value and standard error of mean were calculated for each subject were calculated. The standard errors were calculated to standard deviations. The statistical significant difference between groups were compared by T-test. *p* < 0.0001 was referred as statistically significant.

## Results

### The cardiac ventricular hypertrophy index

In both group of control and microgravity simulation, there is a significant correlation between left ventricular weight and wall thickness measured either as free anterior wall (control n=5, r=0.98; simulation n=5, r=91). There is also a correlation between isolated right ventricular weight and pulmonary outflow tract thickness (control n=5, r=0.89; simulation n=5, r=0.90) in control and simulation group.

In simulation group, but not the control group, the whole heart and left ventricular weight was significantly improved (grey columns in **Figure 1a**, *** *p* < 0.0001). This increasing is the tail-suspension induced fluid redistribution relation over preload in the heart. The left ventricle wet weight to the whole heart ratio was significantly increased after simulation (grey column in **Figure 1b**, *** *p* < 0.0001). A slight weight increasing was observed in simulation right ventricle, but no statistic significant (white column in **Figure 1a**). The right ventricular weight to whole heart weight ratio have no significant difference (white column in **Figure 1b**)

**Figure 1.**
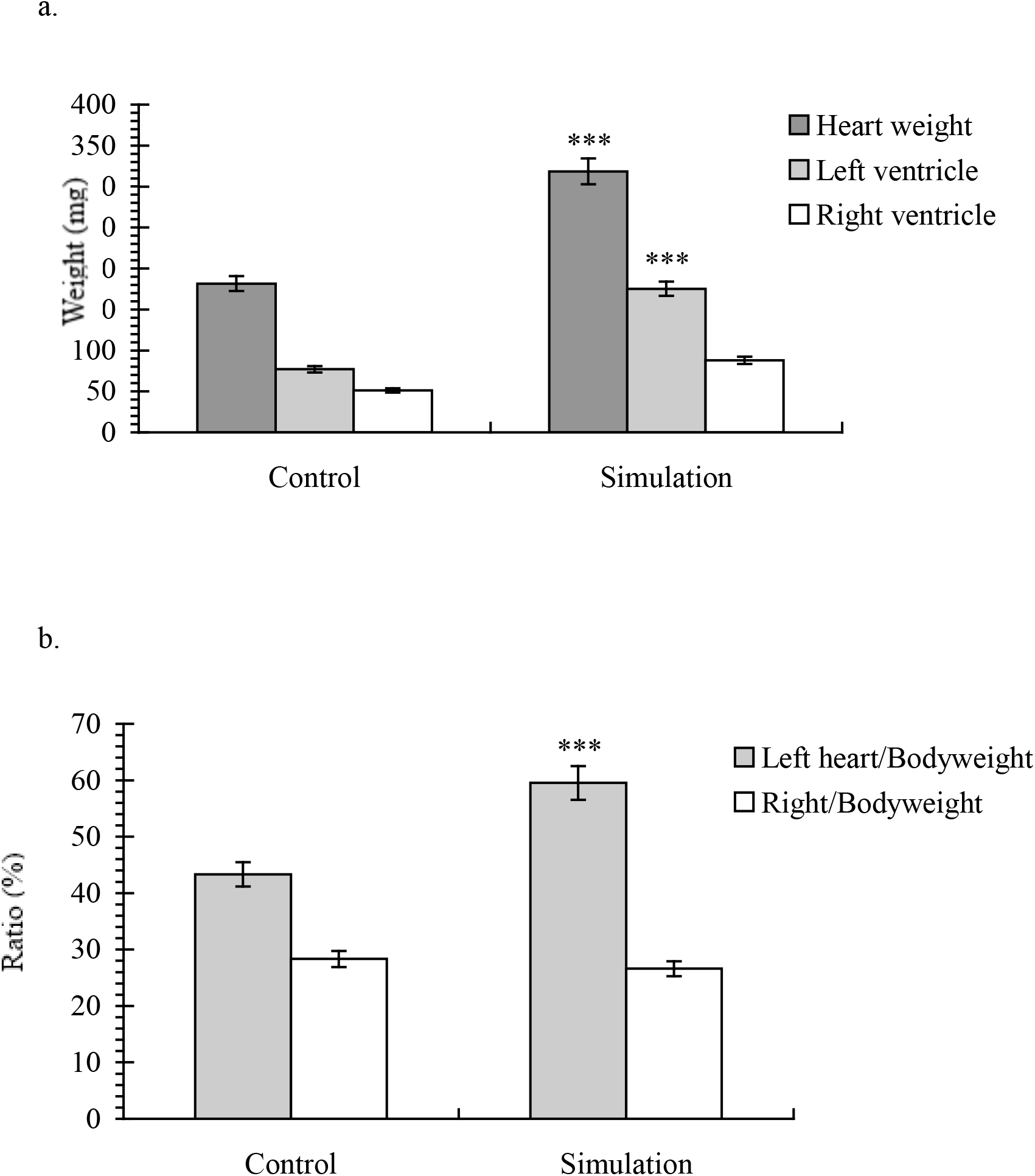
The rat cardiac ventricle myocardium remodeling after 2 weeks’ microgravity simulation.

### The specific STAT3 expression

In the H.E. staining images, there were no significant morphological changes of microgravity simulation cardiomyocytes in right ventricle. The intercalated disc was normal in most area of the simulation right ventricular wall. However, in the vertical section image, in varies area of simulation right ventricle, the cardiomyocyte membrane was not integrity (**Figure 2a**, →). The cardiomyocytes appeared more separated from each other compared with those of the control group (**Figure 2a**, *). of both adjacent cells lined up in the form of corrugated internodes, and there was a depopulation in the side and side connection between both cells (vertical section, *). All of these morphological changes revealed an underlying right ventricular functional dissociation after microgravity simulation.

**Fig. 2.**
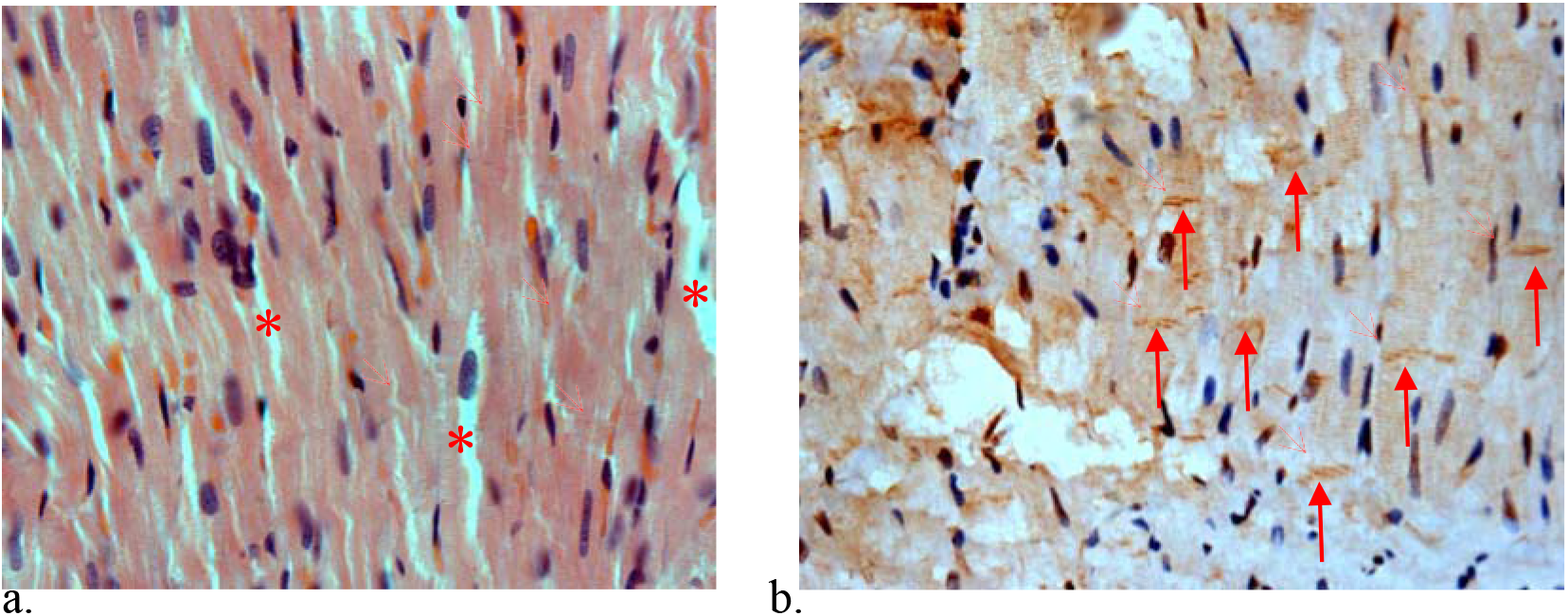
Rat right ventricle myocardium intercalated disc after 2 weeks microgravity simulation.

In simulation right ventricle, cardiac-specific STAT3 signaling were accumulated in the intercalated disc area, presented a kind of slubby expression ((**Figure 2b**,→). The STAT3 accumulation is the primary stage scenario of dysregulation, fibrosis, and decreased cardiac function. STAT3 response the body fluid redistribution, inhibited the restored normal cardiac structure and function in over preload heart remodeling.

## Discussion

In microgravity, the scenario of myocardial growth stimulation is altered. On the systemic level, there are changes in hemodynamic and neuroendocrine regulation that exert indirect effects on the myocardium ^[4]^. In control right ventricle, cardiac muscle fibers are connected to each other through the intercalated disc, which ensures force transmission and transduction between cells and allows the myocardium to function in synchrony. Intercalated disc possesses an intrinsic ability to sense mechanical changes, to achieve a biochemical response that results in molecular and cellular changes in cardiomyocytes. This becomes of particular importance in cardiomyopathies, where the heart is exposed to increased mechanical load and needs to adapt to sustain its contractile function ^[5]^. The mechanosensing and transduction at the intercalated disc can occur mostly through conformational changes in proteins of the adherens junctions complex: a change in the cytoplasmic domain of classical cadherins ^[6]^. In simulation microgravity condition, rat ventricular cardiomyocytes appeared more separated from each other and were slightly smaller in size, exhibiting focal disorganization of muscle fibers and some degenerating cardiomyocytes, of which the nuclei had become pyknotic or disappeared ^[7]^. Our experiment results suggested that simulation microgravity did not increase the significant right ventricle hypertrophy due to the increase of redistribution of the body fluid. However, a part of the right ventricular myocardium caused depopulation of cell interval. In most area of simulation right ventricular wall, the intercalated disc has no significant variation in H.E. staining images. However, the STAT3 signaling protein were abnormal accumulated in intercalated disc in simulation heart, this phenomenon did not appear in control rats.

On the basis of the most recent description of the intercalated disc organization, the functional collagen adhesin gene is considered as a disease of the intercalated disc, are the early molecular events leading to cardiomyocyte degeneration, fibrosis and fibro-fatty substitution. The recent advances in the link between the molecular genetics and pathogenesis of ACM and of the novel role of cardiac intercalated discs. This contribute to the arrhythmogenic cardiomyopathy ^[8]^. In dilated cardiomyopathy variated intercalated discs lead to disturbed contraction and force transduction. The subcellular alterations in sick ventricles intercalated disc proteins were destabilized during cardiac injury in adults ^[9]^. In our experiment, the rearrangement of the intercalated disc was from the effective redistribution of body fluid, thus increasing the blood venous and right ventricle return, further increased the preload in the right ventricle. In such ventricle, cardiac muscle fiber STAT3 protein signal demonstrated a significant accumulation. The SH2 domain of STAT3 has a function to recognize and bind phosphotyrosine residues in the myocardium gp130 receptor signaling chain. The STAT3 positive area demonstrated the STAT3 SH2 domain involved in the intercalated disc remodeling. Furthermore, STAT3 was indicated have the activation through IL-6/IL-11 in cancer-associated fibroblasts promotes tumour development ^[10]^. The effect of STAT3 specific accumulation in remodeling intercalated discs is still not well known, but perhaps the accumulated signaling triggering the fibrosis in over preload right ventricle.

Finally, STAT3 may play a role in inhibiting the replication of covid-19 in host cells. The high expression STAT3 in tail-suspension heart can become a new research model for study covid-19 infection in cardiac muscle fibers.

## Acknowledgements

This research was carried out by the University Student Innovation Challenge promotion project (202011810020).

